# Dynamics of a combined medea-underdominant population transformation system

**DOI:** 10.1101/005512

**Authors:** Chaitanya S. Gokhale, Richard Guy Reeves, Floyd A Reed

## Abstract

**Background:** Transgenic constructs intended to be stably established at high frequencies in wild populations have been demonstrated to “drive” from low frequencies in experimental insect populations. Linking such population transformation constructs to genes which render them unable to transmit pathogens could eventually be used to stop the spread of vector-borne diseases like malaria and dengue.

**Results:** Generally, population transformation constructs with only a single transgenic drive mechanism have been envisioned. Using a theoretical modelling approach we describe the predicted properties of a construct combining autosomal Medea and underdominant population transformation systems. We show that when combined they can exhibit synergistic properties which in broad circumstances surpass those of the single systems.

**Conclusion:** With combined systems, intentional population transformation and its reversal can be achieved readily. Combined constructs also enhance the capacity to geographically restrict transgenic constructs to targeted populations. It is anticipated that these properties are likely to be of particular value in attracting regulatory approval and public acceptance of this novel technology.

## BACKGROUND

Curbing the spread of vector borne diseases such as malaria or dengue is possible by eliminating the transmission capabilities of the insect vectors. One of the many approaches to achieve this is population transformation of vector species. In the most commonly discussed application of population transformation the aim is to introduce transgenes into insect populations which render them refractory to spreading diseases. Usually the technique seeks to use evolutionary principles to establish such transgenes at high frequency in populations through the release of genetically transformed stocks (also called population replacement, [1]). Synthetic disease refractory genes have already been developed for human malaria, dengue fever and avian malaria [2–5]. However, to stably transform insect populations with transgenes that are not selectively advantageous it will be necessary to link refractory transgenes to systems that drive them to high frequency in a population [1, 6–8]. Three transgenic population transformation systems have been shown to be effective in laboratory populations of insects. One is a homing endonuclease based system (HEG), which works by converting heterozygotes to homozygotes [9]. The remaining two systems work by reducing the average fitness of heterozygotes and are: Medea [10, 11] and a bi-allelic form of underdominance [12]. Here we explore theoretically a mono-allelic form of underdominance the implementation of which has to date not been published.

While most studies examine the theoretical properties of transgenic constructs embodying single drive mechanisms [7–10, 13], the observation that “most of them have specific characteristics that make them less than ideal” led Huang *et al.* 2007 [14] to explore combinations. They demonstrated that certain combinations resulted in enhanced properties relative to single systems while others had the opposite effect. Here we take an analogous approach for autosomal Medea and mono-allelic underdominance constructs (not examined in Huang *et al.* 2007[14]). We provide a rigorous and flexible analytical framework to explore salient properties across the entire parameter space. Intuitively, the inclusion within a single transgenic construct of more than one drive mechanism provides a degree of resilience to either mutations in the transgenic construct or to drive-resistance alleles which may exist in target population. While the value of this desirable functional redundancy is not analytically explored here, it does however provides an additional motivation for analyzing the properties of combined systems. Similar to Huang *et al.* [14] the motivation for the analysis presented here comes from the realization that intuitive predictions about combined systems can be misleading and that identifying the parameter space where synergistic enhancements occur can motivate technical developments, including the development of mono-allelic form of underdominance.

We briefly summarise the previously known properties of Medea and mono-allelic underdominant systems separately. Then we look in turn at each of the properties of interest and determine if the combined model performs better than each of the techniques independently. The discussion focuses on the impact of a combined system and provides an assessment of its strengths and weaknesses.

### Medea

Natural **M**aternal **e**ffect **d**ominant **e**mbryonic **a**rrest (Medea) alleles were first discovered in *Tribolium* flour beetles [15] and have also been reported in the mouse [16, 17]. They derive their ability to invade populations by maternally induced lethality of wildtype offspring not inheriting a Medea allele (Figure 1) [18]. Thus the wildtype homozygous offspring of the heterozygous mother die with a certain probability *d*. Despite the mechanism(s) by which natural Medea elements exert their maternal effect remaining unknown, Chen et al. [10] were able to generate a synthetic system (*M edea^myd^*^88^) which mimics their evolutionary properties.

**FIG. 1.**
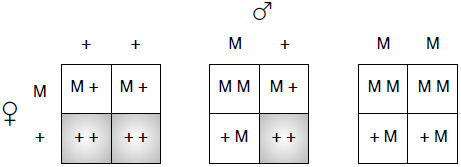
Effect of the Medea allele is seen in offsprings when mothers are heterozygous for Medea. If the mother is a Medea carrier then she deposits a toxin in the oocytes. Only the offspring who have a copy of the Medea allele are rescued. Thus the wildtype homozygous offspring of a heterozygous mother are affected (shaded) and die with a certain probability *d*.

To date, the only published Medea construct (*M edea^myd^*^88^) has been inserted on an autosomal chromosome in *D. melanogaster* [10, 11]. Autosomal Medea insertions unlike sex-chromosome insertions [13] exhibit a high-frequency stable equilibrium when the transgenic construct is associated with any fitness cost (see Figure 2a). As described previously [7, 8, 10, 13, 18] this stable equilibrium results in the persistence of wildtype alleles in populations transformed with autosomal Medea constructs (Figure 2a, though if a linked refractory gene is dominant this is likely to prove unproblematic from the perspective of target disease control).

**FIG. 2.**
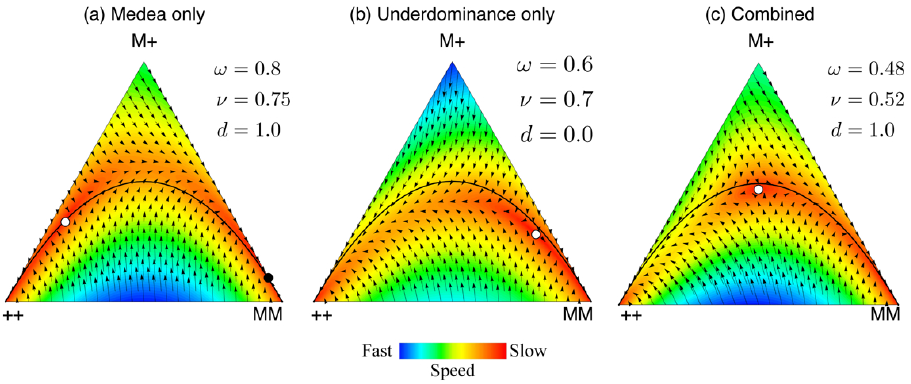
de Finetti diagrams for example parameters. At the vertices the complete population consists of the genotype given by that vertex (++ is for the wildtype homozygote, M+ for the heterozygotes and MM for the transgenic homozygotes). In the interior the population composition is a combination of all the three genotypes with frequencies proportional to the perpendicular distance from the vertex. Unstable equilibrium points are shown as white circles and are always internal within the simplex. Stable equilibrium are shown as black circles and occur on edges (the equilibria which always exist at the ++ and MM corners are not shown). The fitness of the wildtype homozygote is assumed to be 1 and the fitnesses of the other genotypes relative to it are given by *ω* = heterozygotes and *ν* = homozygotes. The lethality effect of the Medea allele is given by the parameter *d*. The three panels describe: **(a)** “Medea only”, an unstable and stable equilibrium occur. These parameters equate to a strong Medea phenotype associated with a significant fitness cost that is substantially dominant. The M allele frequency at the stable threshold is 0.88 and at the unstable threshold is 0.21. **(b)** “Underdominace only”, an unstable equilibrium occurs, always in the right half of the simplex.These parameters equate to weak underdominace with a significant fitness cost in transgenic homozygotes. The unstable threshold frequency of the M allele is 0.8. **(c)** A combined Medea and underdominance system, shows only an unstable equilibrium occurs. We assume multiplicative fitness for *ν* from the values in (a) and (b), The unstable threshold frequency of the M allele is 0.5, which is the ideal threshold for transformation and reversibility (see Appendix). The black line shows the Hardy-Weinberg equilibrium. Note that the system under study easily diverges from the Hardy-Weinberg null model.

### Underdominance

When the heterozygote is less fit than both the possible homozygotes then we have a case of underdominance. However there are only a few examples where alleles at a given locus have been robustly inferred to exhibit underdominance [19]. In a random mating, Hardy-Weinberg population, rarer alleles have larger sojourn times in the heterozygote state, consequently where an underdominant construct is rare it will mostly be in this unfit genotype. Due to the inherently unstable nature of underdominance, if the construct exceeds a threshold value through releases of sufficient homozygotes it is predicted to proceed to fixation within the population (Figure 2b). Intentional underdominant population transformation is inherently reversible where it is realistically possible to release sufficient wildtype individuals to traverse the unstable equilibrium in the lower frequency direction. However, underdominant constructs can be viewed as unappealing when transforming large populations due to the high release numbers required to initiate population transformation (Figure 2b) [20, 21]. The mono-allelic underdominance modeled here describes the situation where there is a transgenic allele at a single autosomal locus (the site of the transgenic construct integration). We have only examined situations where an insert is underdominant in both sexes. A recent publication [12] describing the development of a single locus bi-allelic form of underdominance where there are two functionally distinct transgenic alleles is not applicable to the mono-allelic underdominance analysis described here.

### Medea and underdominance in a single transgenic construct

Here we explore the properties of combining both Medea and underdominance in a single transgenic construct on an autosome. As single locus transgenic underdominance effective in both sexes cannot by definition be configured on sex chromosomes we have modeled both systems on autosomes to permit the most direct comparison between single and combined systems. By combining systems, some properties will be discounted, remain the same or synergistically enhanced. We find a broad parameter space where the applied properties of single systems can be argued to be synergistically enhanced. The principle criteria being: (i) lower transformation threshold, (ii) faster population transformation and (iii) enhanced spatial stability of the transformed population.

## METHODS AND RESULTS

### Genotype fitnesses and expected dynamics

The recursion dynamics are analysed for genotype frequencies as maternal-effect killing violates the Hardy-Weinberg principle. With Medea the action of selection on wildtype homozygotes depends not only on their current state but also on the maternal genotype. Here we have the three genotypes, wildtype homozygous, transgenic homozygous and the heterozygous represented by ++, MM, and M+ respectively. We set the fitness of the wildtype homozygote, ++, to 1. The relative fitnesses of the MM homozygote and the M+ heterozygote are given by *ν* and *ω*. The parameter *d* measures the degree of lethality of homozygous wildtype offspring from Medea carrying mothers, from no Medea effect (*d* = 0) to complete lethality (*d* = 1). Using Table I we calculate the expected frequencies of all three genotypes in the next generation as,

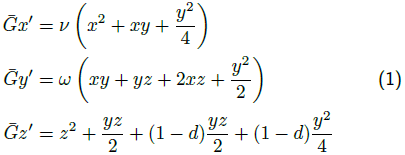

**TABLE I.**
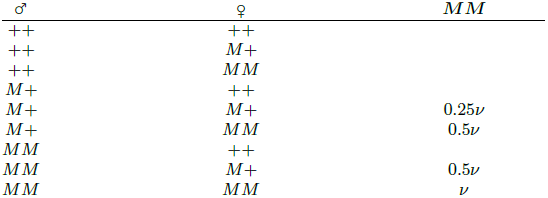
The next generation offspring proportions

where *x*, *y*, and *z* are the frequencies of MM, M+, and ++ respectively in the current generation and *x*′, *y*′, and *z*′ are the expected frequencies in the next generation (in [18] differences in fitness were ascribed to differences in maternal fecundity rather than zygotic genotypes as is done here). The total contribution from all genotypes in the population (i.e., the average fitness) is given by 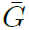. It is the sum of the right hand sides of the set of Eqs. (1) [22]. Another way to view the recursion equations is 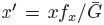, where *f_x_* is the average fitness of the MM genotype [23]. Equating the fitnesses of the three genotypes helps us to solve for the fixed points of this dynamical system (see Appendix). For *d* = 1 there can be an unstable internal equilibrium (Appendix Eq. (A.3)). From the point of view of reversibility it is ideal to have this equilibrium as close as possible to one-half (see Figure 3). This is possible when the fitness values of the heterozygote and the Medea homozygote sum up to unity (see Appendix Eq. (A.5)) as can be seen in Figure 4. The fitnesses of the systems described here are assumed to relate only to the drive mechanism (i.e. without linked refractory genes). In Figure 2 we illustrate how selecting a Medea construct with appropriate parameters results in the combined system having an internal equilibrium closer to the ideal one-half. The release thresholds are determined by the unstable fixed points of the system. As illustrated in Figure 4, combined systems have the potential to be engineered towards an optimal unstable equilibrium value of 0.5 (Eqs. (A.4) and (A.5)). A release threshold substantially smaller than 0.5 would make the construct unappealing from the point of view of reversibility (Figure 3). However, if the size of the target population is large and the capacity to reverse population transformation is not important, then a ‘Medea only’ construct would be the most efficient approach. Solely from the perspective of initiating population transformation, ‘underdominance only’ is disadvantageous as it requires multigenerational release of a large numbers of individuals (albeit smaller than the numbers required for sterile male release programs [24]).

**FIG. 3.**
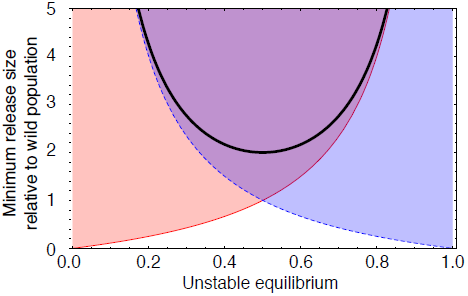
Minimum Release Sizes for Population Transformation. Size of release relative to the wild population is plotted as a function of the unstable equilibrium given by the frequency of the Medea allele 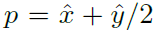. To achieve population transformation the release size must be above the solid red line (*p/*(1 - *p*)). To reverse a transformation the release must be above the dashed blue line ((1-*p*)*/p*). The combined transformation-reversal release sizes are above the thick black line (1*/p*(1 - *p*) - 2), which has a minimum at *p* = 1*/*2.

**FIG. 4.**
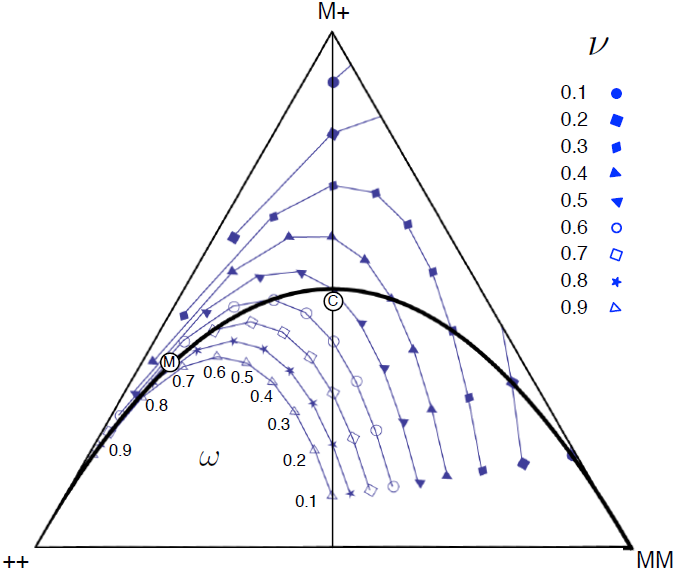
Fitness and its impact on the unstable equilibrium for complete Medea lethality (*d* = 1). The position of the internal unstable equilibrium is illustrated which needs to be traversed for population transformation and for reversal. As the value of *ν* increases the unstable equilibrium moves closer to the all ++ vertex. The different values of *ω* trace a curve which intersects the Hardy-Weinberg equilibrium line at *ν* = *ω*. For underdominance the fixed points are always below the Hardy-Weinberg curve (also see Figure 7). This also graphically demonstrates Eqs. (A.4) and (A.5) i.e. the frequency of the Medea allele is 1*/*2 when *ν* + *ω* = 1 (vertical line, which also represents the ideal with respects to the ease of transformation and its reversal, see Figure 3). Note that when the unstable equilibrium is above the Hardy-Weinberg equilibrium line, there also exists a stable root on the M+ – MM edge given by 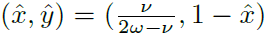. Disks indicate the positions of results plotted in Figure 2 for the ‘Medea only’ system (M) and the combined system (C).

It is generally appreciated that once releases commence, population transformation should occur as rapidly as possible and proceed to complete fixation. This minimizes the possibility of selection for insects resistant to the transformed construct. Furthermore the pathogen itself could evolve mechanisms to evade the effects of the linked refractory genes. A rapid and complete fixation of the transgenic construct and elimination of the pathogen minimizes both possibilities (neither of which are explicitly modeled here). Clearly, releasing as many individuals as is feasible is an effective way to speed population transformation [21, 25]. We show that the time taken to achieve population transformation can also be reduced by combining two systems, where even very weak Medea (*d ≤* 0.2) has a large impact on the speed of transformation. (see Figure 5 and 0.65 starting frequency). The acceleration can also occur during reversal of population transformation. Knowledge of this effect will permit the design of efficient release strategies for both the initiation of population transformation and for its rapid reversal.

**FIG. 5.**
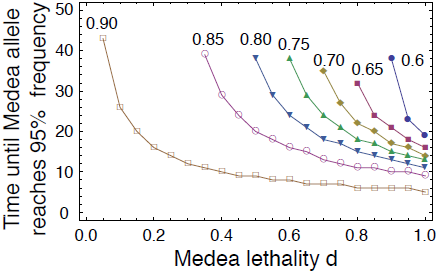
Numerical solutions for critical times starting at different initial frequencies of the MM genotype. With the parameter values for the combined system (*ω* = 0.48*, ν* = 0.52, Figure 2 C) we begin on the ++ − MM edge at different frequencies. The time required to reach MM frequencies *>* 0.95 are plotted as the critical times. Starting with the frequency of MM genotypes of 0.6 (circles) only if *d ≥* 0.85 the system moves to the MM vertex. As the Medea lethality increases the all MM vertex can be reached by starting at lower frequencies of MM genotype. Starting at already high frequencies (0.9, open squares) the time to reach fixation quickly drops to the levels which are almost the same as that of complete Medea lethality. (Initial MM frequencies 0.6 (circles), 0.65 (squares), 0.7 (diamonds), 0.75 (triangles), 0.8 (inverted triangles)). For the recursions, Eqs. (1) were employed.

### Population structure dynamics

We consider a simple two-deme model of population structure, where two populations of large and equal size are coupled by a symmetrical fraction of migrants *m* between the populations in each generation. Considering asymmetries in population sizes, migration needs to be dealt with separately, as in [26]. Also migration dynamics with an explicitly set spatial system has been recently assessed [27] (albeit not for a combined system). In population *i* the expected genotype frequency of genotype *k* after migration is 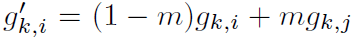, where *g_k,i_* is the frequency of the *k^th^* genotype in population *i* and *g_k,j_* is the *k^th^* genotype frequency in population *j*. These adjusted genotype frequencies can then be substituted into Eqs. (1).

We initialize the two populations where the Medea allele is almost fixed in one and almost lost in the other. The recursions were performed for different migration rates, slowly incremented in units of 10^−3^. The migration rate where the difference in allele frequencies between the two populations fell below 1% (thus assuming populations to have reached an equilibrium), was recorded as the critical migration rate. At lower than critical migration rates the combined systems will not spread far from a successfully transformed zone, and will be resistant to loss by immigration. We evaluated the critical migration rate which allow the transformation of a local population stably (Figure 6). For a varying heterozygote fitness *ω* ranging from 0.01 to 0.95 we consistently see that having a Medea construct provides more geographical stability as compared to a system without Medea even as we move from a system with directional selection against Medea to an underdominant system. Figure 6 shows that a combined system has a higher geographic stability in terms of limiting the unintentional transformation of adjacent wildtype populations for a wide range of values of transgenic homozygote fitness *ν* (‘Medea only’ exhibits limited geographic control if fitness costs of being transgenic are high [8]). Interestingly, in combined systems geographic stability does not increase monotonically with respects to *v*. This can result in maximal geographic stability for combined systems at intermediate values of *ν* (Figure 6). The levels of sustained migration, which maintain geographic stability, can be surprisingly high and of an order expected between highly interconnected demes rather than between isolated populations [28]. In addition to the obvious regulatory benefits of robust geographic stability, this property can be exploited to limit the number of transgenic individuals in which unintended mutational events can occur or to lower the probability that pathogens evolve resistance to linked refractory genes.

**FIG. 6.**
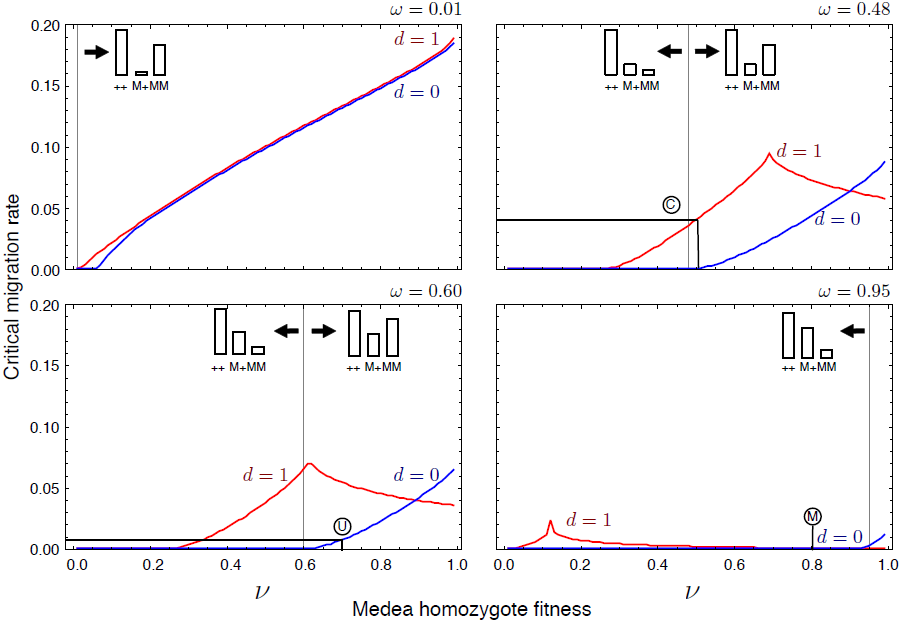
Critical migration rates allowing stable local transformations over a range of genotype and Medea parameter configurations. Using the recursion equations Eq. (1) with modifications as described in the “Population structure dynamics” section we explore the pattern when there is no Medea effect (*d* = 0) and complete Medea lethality (*d* = 1). For different values of the heterozygote fitness (*ω*) we explore the genotype configurations going from directional selection to underdominance. The transition in the fitness structure between these two states is indicated within the plots using token bar charts (illustrated graphically within the plots). Over a wide range of parameter space the combined construct exhibits substantially higher critical migration thresholds than ‘underdominance only’. Interestingly the *d* = 1 dynamics are not monotonic. The figure illustrate how migrational stability can be enhanced, even with a reduction in fitness of the genetically modified homozygote. Disks indicate the positions of results plotted in Figure 2 for the ‘underdominance only’ system (U), the combined system (C) and ‘Medea only’ (M). Comparing the combined system with with ‘Medea only’ system we see that not only Medea but underdominance also is necessary to get the desired migrational stability in experimental systems.

## DISCUSSION AND CONCLUSION

In the theoretical analysis of combined population transformation systems Huang *et al.* [14] considered the combination of a transgenic two-locus form of underdominance (termed engineered underdominance [29] with two other natural phenomena (*Wolbachia* and sex-linked meiotic drive). Both *Wolbachia* and sex-linked meiotic drive were demonstrated to have the potential to significantly impact the feasibility and dynamics of population transformation in both positive and negative ways. It was clearly shown that intuitive expectations of combined systems could be misleading and that mathematical modeling was essential in identifying potentially useful combinations and parameter values (most notably those relating to genotypic fitness). An excellent example is the Huang *et al.* [14] theoretical analysis of the two-locus form of engineered underdominance which has been only recently realised [12].

Here we have followed an analogous approach to explore the properties of combining two currently developed transgenic drive systems within a single autosomal construct. The underdominant and Medea systems are assumed to be physically interspersed in a manner that maximizes the probability that they remain linked (e.g. in a configuration analogous to that shown in Figure 2 [10]). The described modeling framework has allowed us to identify a broad parameter space where combined systems can in some circumstances outperform single systems in terms of (i) optimizing release thresholds (Figures 3, 4, 7) (ii) increasing the speed of population transformation and (Figure 5) (iii) enhancing the geographic stability of population transformation (Figure 6). In addition, the reliance on two distinct mechanisms for population transformation could reduce the probability that resistance to the transgenic construct arises in the target insects. If however, long term selective pressures within successfully transformed target populations would result in the loss of the underdominance mechanism, this essentially leaves a ‘Medea only’ construct at high frequency. This ‘Medea only’ construct would be impractical to remove (unless it was associated with a high fitness cost) and could spread to adjacent populations. Conversely, loss of the Medea mechanism from a combined construct has a considerably smaller impact on reversibility and stability (Figures 2 and 6). Recognizing that the loss of Medea is preferable to loss of underdominance, it would be prudent to engineer underdominance which is more mutationally stable than Medea (duplicating the underdominant mechanism would be one simple strategy). It is also noteworthy that many of the synergistic enhancements ascribed to combined systems are to a significant extent shared by Medea constructs inserted on sex-chromosomes [13]. Consequently, depending on the empirical properties of autosomal versus sex-chromosome inserts the relative merits of both approaches would warrant evaluation within the specific objectives of a given program.

**FIG. 7.**
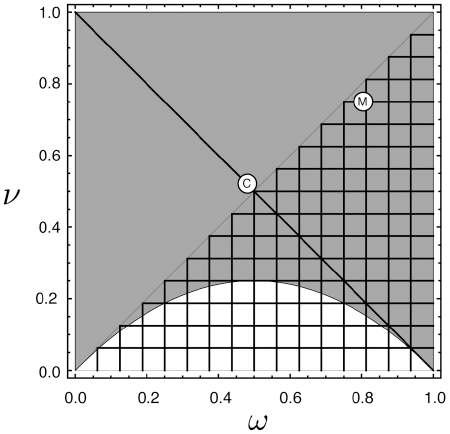
The configuration of the stable and unstable equilibrium in the phase space for **d** = **1**. The high-frequency unstable equilibrium and stable equilibrium were determined numerically for *d* = 1 over a range of fitness values. In the shaded region an unstable equilibrium exists within the interior of the simplex. In the meshed region a stable equilibrium exists on the M+ to MM edge of the simplex The non-mesh region corresponds to underdominance. The dark diagonal line denotes an ideal unstable threshold in terms of ease of populations transformation and reversal (*x*^ + *y*^*/*2 = 1*/*2) (see Appendix). Disks indicate the positions of results plotted in Figure 2 for the ‘Medea only’ system (M) and the combined system (C).

It has been assumed throughout that fitness costs are directly associated with the drive mechanism or mechanisms in a transgenic construct, however it is also likely that additional costs will also be associated with anti-Plasmodial or anti-viral genes included as part of a working construct. The analytical framework described here will permit the prediction of the properties of combined systems loaded with such disease refectory genes. The fitness cost of refractory genes has in some, but not all, circumstances been estimated to be quite high [30]. Consequently the illustrative parameters values used in Figure 2 may represent plausible values for ‘loaded’ constructs (though the framework presented here allows exploration of the entire range of parameters). The immediate practical use of this method could help protect *D. melanogaster* from an unintended species wide Medea transformation if combined with underdominance for testing in the lab. The most likely application of population transformation is in species of the genera *Anopheles* and *Aedes* which act as devastating disease vectors [7]. Within these genera there are significant differences in dispersal capacities estimated at various locations, in some instances individuals migrate hundreds of meters over their lifetime [31]. Consequently, the capacity to restrict transgenic constructs to particular populations is likely to be considered of high value. Various configurations of underdominance have been proposed as representing the most likely system to maintain geographic stability [12, 29]. Geographic stability is generally achieved by maximizing the fitness of transgenic homozygotes fitness *ν*. However where this is not possible due to cost arising from the underdominant drive mechanism or of refractory genes, our analysis indicates that maximal geographic stability can be achieved by combining systems for intermediate values of *ν* (Figure 6). Exploitation of this phenomena, in addition to the value of functional redundancy in drive mechanisms, could provide a valuable practical incentive to explore combined drive systems experimentally.

## AUTHORS’ CONTRIBUTIONS

CSG, RGR and FAR conceived the project. All authors developed the model. CSG and FAR performed simulations. All authors analysed the results and wrote the manuscript. All authors read and approved the final manuscript.

## ACKNOWLEDGEMENTS

We thank P. M. Altrock, J. F. Baines, J. Denton, and A. Traulsen, for discussions and V. L. Reed for comments on the manuscript. CSG is supported by the Emmy-Noether program of the Deutsche Forschungsgemeinschaft.

RGR is supported by grant RE-3062/2-1 of the Deutsche Forschungsgemeinschaft. FAR is supported by the College of Natural Sciences, University of Hawai‘i at Mānoa, by the Max Planck Society, and by a grant from the Hawai‘i Community Foundation.

### APPENDIX

#### Average genotype fitnesses and calculating the equilibria

The frequencies of the genotypes in the next generation are given by *x*′, y′ and *z*′. In equilibrium we have *x*′ = *x*, *y*′ = *y* and *z*′ = *z*. However the expressions for the next generation frequencies are rational functions given by, 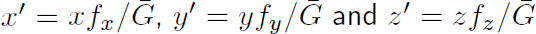 where the fitnesses of the genotypes are given by,

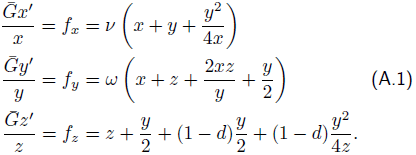

Now in equilibrium the frequencies of the genotypes do not change over generations but it is a consequence of their average fitnesses being the same. Hence we can deduce the equilibria of the system just be equating the average fitnesses. This is just another way of writing *x*′ = *x*, *y*′ = *y* and *z*′ = *z*, which reduces to 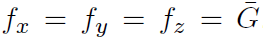. Considering the average fitness of the genotypes in a pairwise fashion, two genotypes are neither increasing or decreasing relative to each other if their average fitnesses are equal, e.g., *f_x_* = *f_y_*. This argument is obvious when we view the system in continuous time. While the recursion equations predict the dynamics of the system in the next time step, one at a time, we can explore the complete dynamics by analysing the analogous difierential equations given by,

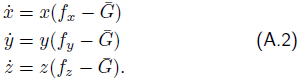

where the time derivative of a variable is given by *ẋ* = *dx = dt* and so forth for *y* and *z*. From the form of these difierential equations the equilibrial solutions are evident, either when the frequencies are zero (vertices of the simplexes in Figure 2) or when the bracketed terms are zero. Since the genotype frequencies sum up to 1, we can solve for just two frequencies. The solutions obtained though are complicated expressions with a possibility of imaginary roots.

Assuming complete Medea lethality (*d* = 1), the equilibrium of the system is given by,

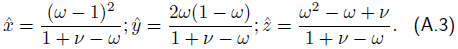

When it exists 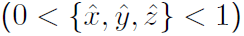 then it is always unstable. Of particular interest is the case where the Medea allele frequency (given by 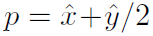) at the unstable equilibrium is 0.5,

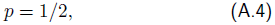

as this is ideal from the point of view of reversibility (see Figure 3). In order to cross an unstable equilibrium threshold, to ultimately transform a population, releases have to be made of a minimum size of *p*/(1*–p*) relative to the wild population size. To cross this boundary and then recross it (i.e. if we wish to reverse a completely transformed population) requires two releases with a minimum combined size of 1/*p*(1 *– p*) *–* 2. This function approaches positive infinity at *p* = 0 and *p* = 1 and has a minimum at *p* = 1/2 with the release ratio being twice that of the wild population (Figure 3). Thus, an unstable threshold of *p* = 1/2 is ideal from the perspective of population transformation and reversibility and is still much lower than release sizes used in successful applications of the sterile insect technique. Substituting the equilibrium values in Eqs. (A.3) into Eq. (A.4) gives

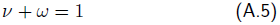

at *d* = 1 (Figure 7). For *t <* 1, there may exist two internal equilibria, the lower allele frequency one is unstable and the higher frequency one is stable, given by

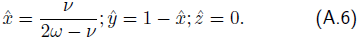

However, in case of underdominance if Eq. (A.5) holds then only the unstable internal equilibrium exists at *p* = 1/2. This is graphically illustrated in Figure 7.

